## supplementary file for "CoRNeA: A pipeline to decrypt the inter protein interfaces from amino acid sequence information"

***Corresponding Author**:

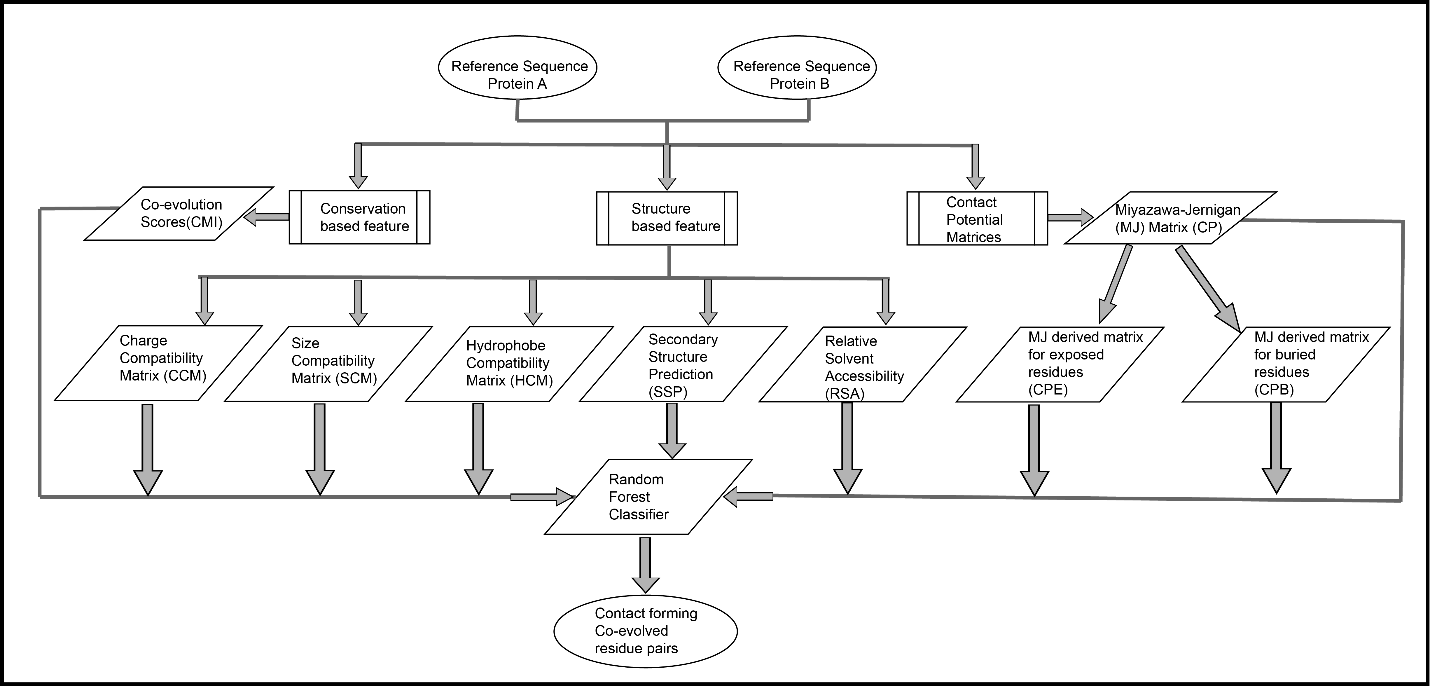

**Figure S1: Flowchart depicting the feature generation for predicting pair of protein-protein interaction interface residues**

**Table S1: Numeric Coding for amino acids used for co-evolution score calculations**

| **Amino Acid** | **Numeric Coding** |
| --- | --- |
| V (Valine) | 1 |
| I (Isoleucine) | 2 |
| L (Leucine) | 3 |
| M (Methionine) | 4 |
| F (Phenylalanine) | 5 |
| W (Tryptophan) | 6 |
| Y (Tyrosine) | 7 |
| S (Serine) | 8 |
| T (Threonine) | 9 |
| N (Asparagine) | 10 |
| Q (Glutamine) | 11 |
| H (Histidine) | 12 |
| K (Lysine) | 13 |
| R (Arginine) | 14 |
| D (Aspartic Acid) | 15 |
| E (Glutamic acid) | 16 |
| A (Alanine) | 17 |
| G (Glycine) | 18 |
| P (Proline) | 19 |
| C (Cysteine) | 20 |
| - (Gap) | 21 |
| X (Non-Standard Amino Acid) | 22 |

**Table S2: Comparison of known methods for PPI interface prediction with the new hybrid method**

| **Interface residues (PISA)** | | | **Various algorithms for finding contacts** | | | | |
| --- | --- | --- | --- | --- | --- | --- | --- |
| **Nup107** | **Nup133** | **Distance(Å)** | **MI**  **(2.03)** | **DCA**  **(0.158)** | **Evfold**  **(0.155)** | **SCA**  **(3.86)** | **New Method (CMI)**  **(1.00)** |
| **D 879** | **T 696** | 3.37 | 0.4285 | 0.0022 | 0.0052 | 0.618 | **0.804** |
| **S 822** | **K 975** | 2.78 | 0.2379 | 0.0009 | 0.0023 | 0.1607 | **0.591** |
| **E 884** | **K 975** | 2.69 | 0.2379 | 0.0001 | 0.0021 | 0.339 | **0.524** |
| **D 917** | **K 966** | 2.53 | 0.0104 | 0.0005 | 0.0013 | 0.192 | **0.642** |
| **Y 921** | **K 966** | 3.37 | 0.225 | 0.0008 | 0.003 | 0.616 | **0.364** |
| **E 922** | **R 962** | 3.18 | 0.7898 | 0.0015 | 0.002 | 0.742 | **0.342** |
| **K 894** | **D 982** | 3.82 | 0.354 | 0.005 | 0.0005 | 0.223 | **0.371** |
| **R 898** | **A 980** | 3.28 | 0.179 | 0.001 | 0.0025 | 0.039 | **0.233** |
| **Q 902** | **Q 944** | 3.35 | 0.8474 | 0.002 | 0.001 | 1.46 | **0.159** |

The interface residues for a test case as predicted by PISA. The value under the name of the method represents the highest score calculated by the algorithm. MI: Mutual information, DCA: Direct Coupling Analysis, SCA: Statistical Coupling Analysis.

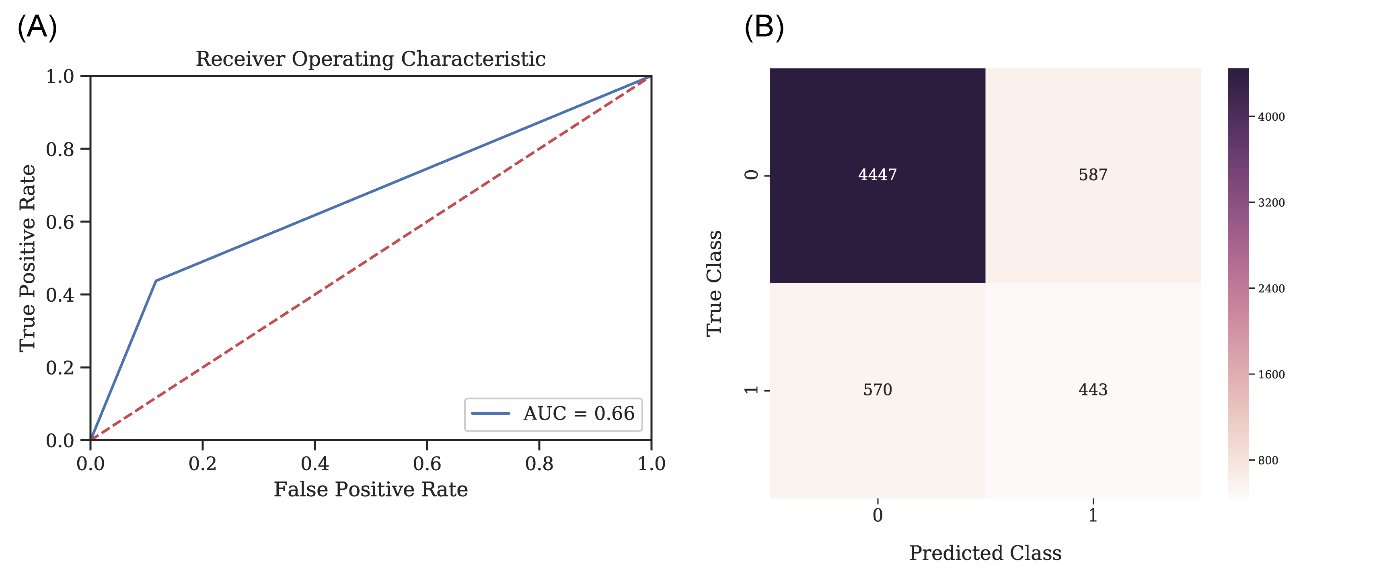

**Figure S2: Statistics for the Random Forest Classifier Model for predicting contact forming residue pairs without environmental features. (**A) Receiver-operator curve (ROC) depicting Area under the curve (AUC) as 0.66 when the model is tested on the 75:25 data split. (B) Confusion matrix for the tested model on 75:25 data split with a final accuracy of 80%

**Table S3: Comparison of evaluation statistics, with and without environmental features.**

|  | **Class** | **Precision** | **Recall** | **F1-score** |
| --- | --- | --- | --- | --- |
| **Without Environmental Features** | **0** | **0.89** | **0.88** | **0.88** |
|  | **1** | **0.43** | **0.44** | **0.43** |
|  | **Weighted Avg** | **0.81** | **0.81** | **0.81** |
| **With Environmental Features** | **0** | **0.92** | **0.91** | **0.91** |
|  | **1** | **0.56** | **0.59** | **0.58** |
|  | **Weighted Avg** | **0.86** | **0.85** | **0.86** |

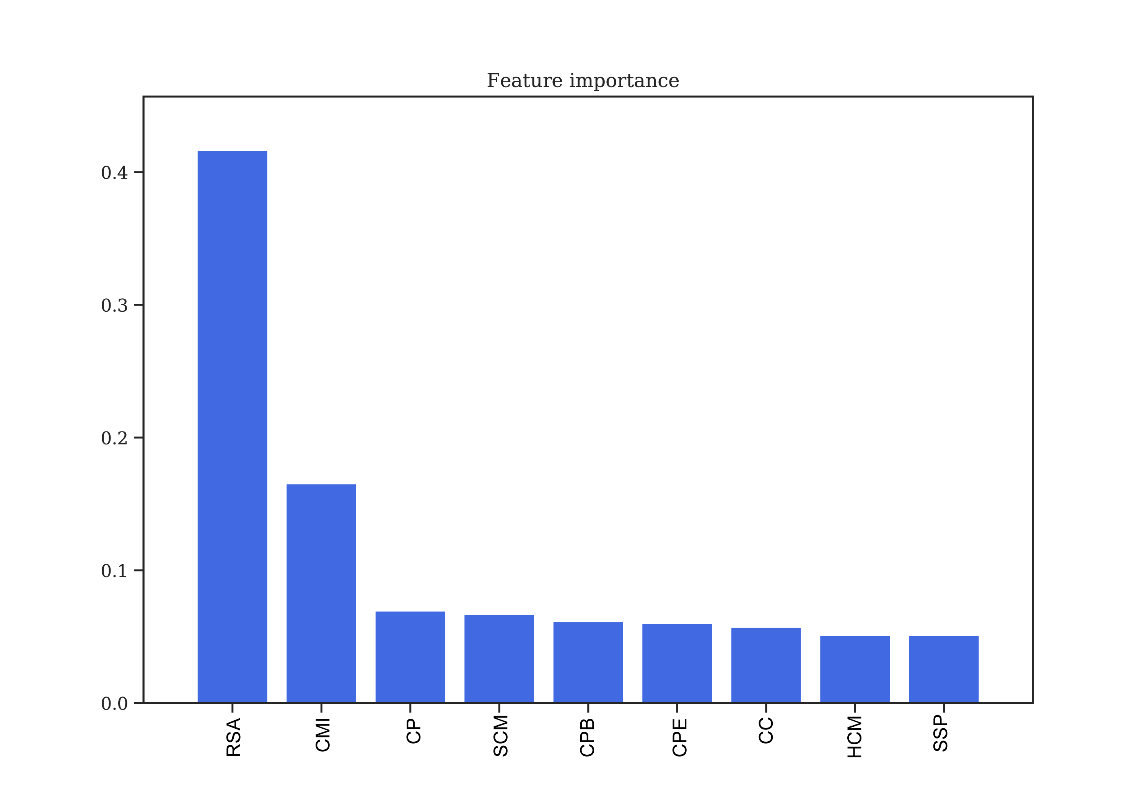

**Figure S3: Feature Importance obtained from Random Forest Classifier without environmental features.**

Relative Solvent Accessibility (RSA) and Co-evolution Scores (CMI) as two of the most important features in training the model. **RSA:** Relative Solvent Accessibility. **CMI:** Conditional Mutual Information. **CP:** Contact Potential. **SCM:** Structure Compatibility Matrix. **CPB:** Contact Potential for Buried residues. **CPE:** Contact Potential for Exposed residues. **CC:** Charge Compatibility. **HCM:** Hydropathy Compatibility Matrix. **SSP:** Secondary Structure Prediction.

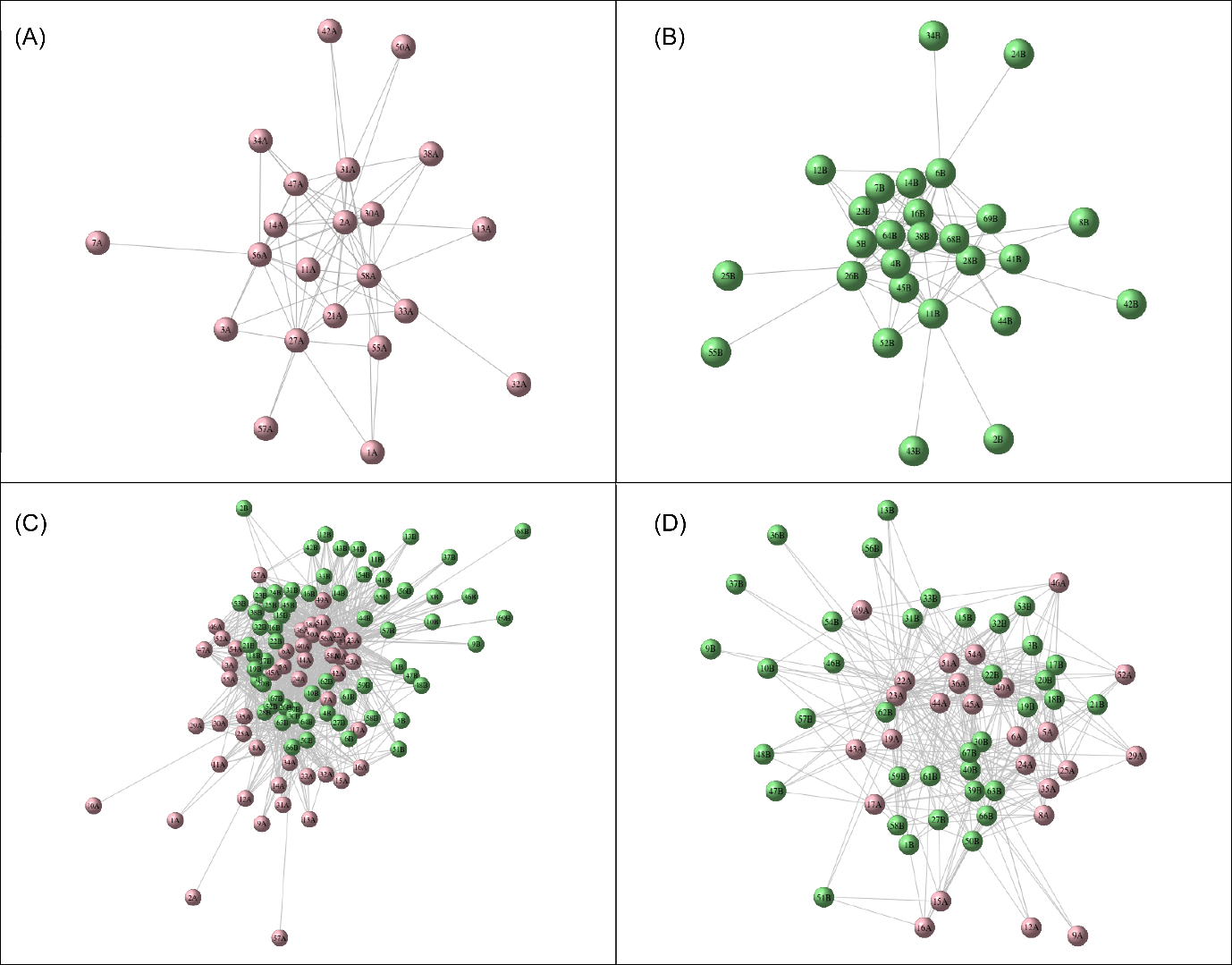

**Figure S4: Network analysis for PDB ID 1GCQ.** (A) Intra-protein network for Chain A/B of 1GCQ obtained from top 5% co-evolving intra residue pairs. (B) Intra-protein network for Chain C of 1GCQ obtained from top 5% co-evolving intra residue pairs. (C) Inter-protein network for 1GCQ obtained from random forest classifier. (D) Inter-protein network for 1GCQ after removing intra-protein network nodes and all nodes having relative solvent accessibility as 0.

**Table S4: Pairwise true contacts predicted for PDB ID 1GCQ Chain A with Chain C and Chain B with Chain C within a distance cutoff of 10 Å.**

| **Residue number**  **(Chain A)** | **Residue number**  **(Chain C)** | **Distance (Å)** | **Residue number**  **(Chain B)** | **Residue number**  **(Chain C)** | **Distance (Å)** |
| --- | --- | --- | --- | --- | --- |
| **208** | **609** | **3** | **179** | **652** | **3.3** |
| **208** | **608** | **3.3** | **165** | **657** | **3.6** |
| **209** | **610** | **3.5** | **179** | **637** | **4** |
| **192** | **611** | **3.6** | **165** | **656** | **5.3** |
| **208** | **611** | **3.6** | 211 | 629 | 5.9 |
| **193** | **610** | **4** | 179 | 653 | 6.6 |
| **193** | **611** | **4** | 165 | 653 | 7.25 |
| **208** | **612** | **4.3** | 179 | 651 | 7.7 |
| **192** | **612** | **4.4** | 179 | 636 | 8 |
| **165** | **608** | **4.8** | 179 | 656 | 8 |
| **209** | **611** | **4.9** | 179 | 657 | 8 |
| 208 | 610 | 5.2 | 209 | 612 | 8.3 |
| 193 | 612 | 5.6 | 163 | 657 | 8.3 |
| 206 | 612 | 6 | 179 | 630 | 8.7 |
| 193 | 609 | 7.3 | 182 | 630 | 8.8 |
| 208 | 607 | 7.7 | 179 | 627 | 9 |
| 192 | 609 | 7.7 | 180 | 637 | 9 |
| 166 | 653 | 7.8 | 208 | 593 | 9.3 |
| 179 | 607 | 8.5 | 211 | 593 | 9.3 |
| 165 | 609 | 8.7 | 179 | 629 | 9.5 |
| 193 | 608 | 8.8 | 179 | 600 | 10 |
| 165 | 610 | 8.9 | 180 | 630 | 10 |
| 209 | 653 | 9.3 | 211 | 652 | 10 |
| 192 | 608 | 9.6 | 192 | 657 | 10 |
| 165 | 651 | 9.6 | 211 | 657 | 10 |
| 179 | 608 | 9.8 |  |  |  |
| 174 | 612 | 10 |  |  |  |

**Table S5: Confusion Matrix statistics for PDB ID 1GCQ before and after network analysis**

| **Before Network Analysis** | **0**  **True Class 1** | **True Negatives= 2954** | **False Positives = 967** |
| --- | --- | --- | --- |
|  |  | **False Negatives= 56** | **True Positives= 25** |
|  |  | **0 Predicted Class 1** | |
| **After Network Analysis** | **0**  **True Class 1** | **True Negatives= 3575** | **False Positives = 319** |
|  |  | **False Negatives= 56** | **True Positives= 52** |
|  |  | **0 Predicted Class 1** | |

**Table S6: Comparison of predictions from CoRNeA with BIPSPI**

|  | **Method** | **Expected no of residues within 10Å** | **Number of True positives with probability more than 0.5** | **Number of False Positives** |
| --- | --- | --- | --- | --- |
| **PDB ID: 1GCQ** | **BIPSPI** | **108** | **0** | **N/A** |
|  | **CoRNeA** |  | **52** | **56** |
| **PDB ID: 5YVT** | **BIPSPI** | **164** | **24** | **1210** |
|  | **CoRNeA** |  | **24** | **968** |

The numbers depicted for CoRNeA are post network analysis. For 1GCQ the total number of expected contacts and true positives are for both chain combinations i.e. Chain A and C; Chain B and C
